# The dispensability of 14-3-3 proteins for the regulation of human cardiac sodium channel Na_v_1.5

**DOI:** 10.1101/2022.10.26.513875

**Authors:** Oksana Iamshanova, Anne-Flore Hämmerli, Elise Ramaye, Arbresh Seljmani, Daniela Ross-Kaschitza, Noëlia Schärz, Maria Essers, Sabrina Guichard, Jean-Sébastien Rougier, Hugues Abriel

## Abstract

**Background:** 14-3-3 proteins are ubiquitous proteins that play a role in cardiac physiology (*e.g*., metabolism, development, and cell cycle). Furthermore, 14-3-3 proteins were proposed to regulate the electrical function of the heart by interacting with several cardiac ion channels, including the voltage-gated sodium channel Na_v_1.5. Given the many cardiac arrhythmias associated with Na_v_1.5 dysfunction, understanding its regulation by the protein partners is crucial.

**Aims:** In this study, we aimed to investigate the role of 14-3-3 proteins in the regulation of the human cardiac sodium channel Na_v_1.5.

**Methods and Results:** Amongst the seven 14-3-3 isoforms, only 14-3-3η (encoded by *YWHAH* gene) weakly co-immunoprecipitated with Na_v_1.5 when heterologously co-expressed in tsA201 cells. Total and cell surface expression of Na_v_1.5 was however not modified by 14-3-3η overexpression or inhibition with difopein, and 14-3-3η did not affect physical interaction between Na_v_1.5 α-α subunits. The current-voltage relationship and the amplitude of Na_v_1.5-mediated sodium peak current density were also not changed.

**Conclusions:** Our findings illustrate that the direct implication of 14-3-3 proteins in regulating Na_v_1.5 is not evident in a transformed human kidney cell line tsA201.

**Summary:** This work shows that only 14-3-3η, exhibits weak/transient interaction with Na_v_1.5, and does not modify its total protein expression, plasmalemmal trafficking, and basal biophysical properties of the whole-cell current. Furthermore, inhibition of endogenous 14-3-3/ligand interactions with difopein does not affect the dimerization of Na_v_1.5. Therefore, 14-3-3 proteins are suggested to be dispensable for the Na_v_1.5 regulation in a heterologous expression system.

## Introduction

In the heart voltage-gated sodium channel, Na_v_1.5 (encoded by *SCN5A* gene), is responsible for the genesis and propagation of the action potential in the contractile myocardium. The function of Na_v_1.5 largely relies on the interaction with its protein partners that regulate proper localization, turnover, and biophysical properties of the channel (Dong et al., 2020). Dysfunction of Na_v_1.5 is associated with various lethal diseases such as cardiac arrhythmias and cardiomyopathy. Therefore, understanding the relationship between Na_v_1.5 and its regulatory protein partners may uncover novel therapeutic strategies for patients suffering from *SCN5A*-related diseases.

One of the protein partners of Na_v_1.5 was reported to be 14-3-3 proteins, which are a family of conserved regulatory molecules expressed in all eukaryotic cells. 14-3-3 can bind target proteins, modulating protein-protein interactions and the activity of their ligands due to inducing structural rearrangements, masking or unmasking of the functional sites, and changing the ligand’s intracellular localization (Obsil and Obsilova, 2011). *YWHAB*, *YWHAG*, *YWHAE*, *SFN*, *YWHAH*, *YWHAQ*, and *YWHAZ* genes encode the mammalian 14-3-3 isoforms β/α, γ, ε, σ, η, θ, and ζ/δ, respectively, where α and δ represent the phosphorylated forms. All 14-3-3 isoforms exhibit a high level of structural homology and consist of 9 antiparallel α-helices arranged in an L-shape (Obsil and Obsilova, 2011; Yang et al., 2006). Through their N-terminal region, 14-3-3 proteins form homo- and heterodimers (Yang et al., 2006). Each ∼30-kDa monomer binds to a variety of the target proteins at the consensus sequences R(S/X)XpSXP, RXXXpSXP, and pS/pTX1–2-COOH of the phosphorylated motifs (Thompson and Goldspink, 2022). The best-studied examples of the unphosphorylated targets are P*seudomonas aeruginosa* virulence factor exoenzyme S and R18, a peptide identified from a phage display library (Henriksson et al., 2002; Wang et al., 1999). R18 is widely used as a potent antagonist of 14-3-3 protein/ligand interactions due to its high affinity with all 14-3-3 isoforms (*K*_d_ = 80 nM) followed by occlusion of their ligand-binding groove (Wang et al., 1999). Accordingly, dimer of R18, difopein, is also a competitive non-isoform specific inhibitor of 14-3-3 protein/ligand interactions (Masters and Fu, 2001).

14-3-3 proteins target a wide variety of ligands and regulate many cellular processes (Obsil and Obsilova, 2011). Accordingly, 14-3-3 proteins were shown to be involved in cardiac physiology, including cardiomyocyte development, cell cycle, and Ca^2+^ signaling (Gittenberger-de Groot et al., 2016; Kosaka et al., 2012; Martens et al., 2021; Menzel et al., 2020). Importantly, 14-3-3 was proposed to play a role in cardiac electrophysiology due to their interactions with cardiac ion channels, exchangers, and pumps (Thompson and Goldspink, 2022). Specifically, the α-subunit of Na_v_1.5 channel was shown to coimmunoprecipitate with 14-3-3 proteins in mouse heart tissue as well as in heterologous expression systems (Allouis et al., 2006). Although 14-3-3η, -θ, and -ζ were identified by yeast 2-hybrid screen to directly interact with Na_v_1.5, only 14-3-3η was investigated for functional consequences on the channel activity (Allouis et al., 2006). In the presence of exogenous 14-3-3η, the density of sodium current and activation curve were unchanged, while the inactivation curve was negatively shifted (Allouis et al., 2006). Based on this data, computer simulations suggested proarrhythmic effects of 14-3-3η (Allouis et al., 2006). Nevertheless, transgenic mice expressing a cardiac-specific loss-of-function 14-3-3η mutant exhibited normal cardiac morphology, cardiomyocyte appearance, and ventricular systolic function under basal conditions, while nothing was reported about its cardiac electrical activity (Xing et al., 2000). Interestingly, immunostaining analysis revealed colocalization between exogenous 14-3-3η and Na_v_1.5 in the intercalated discs of adult rabbit cardiomyocytes, suggesting a role in clusterization (Allouis et al., 2006). In line with these findings, 14-3-3 proteins were reported to mediate the coupled gating of Na_v_1.5 (Clatot et al., 2017). Although two 14-3-3 binding sites were identified in the intracellular loop between domains I and II of Na_v_1.5 by co-immunoprecipitation analysis, it was unclear which isoform mediated the channel gating properties through its direct interaction (Clatot et al., 2017). Therefore, our goal was to investigate the direct role of all 14-3-3 isoforms on Na_v_1.5 in a heterologous expression system. Specifically, we addressed their interaction, expression, trafficking, and sodium current.

## Materials and Methods

### cDNA constructs and cloning

Template cDNA constructs and their derivatives are listed in Table 1.

**Table 1.**
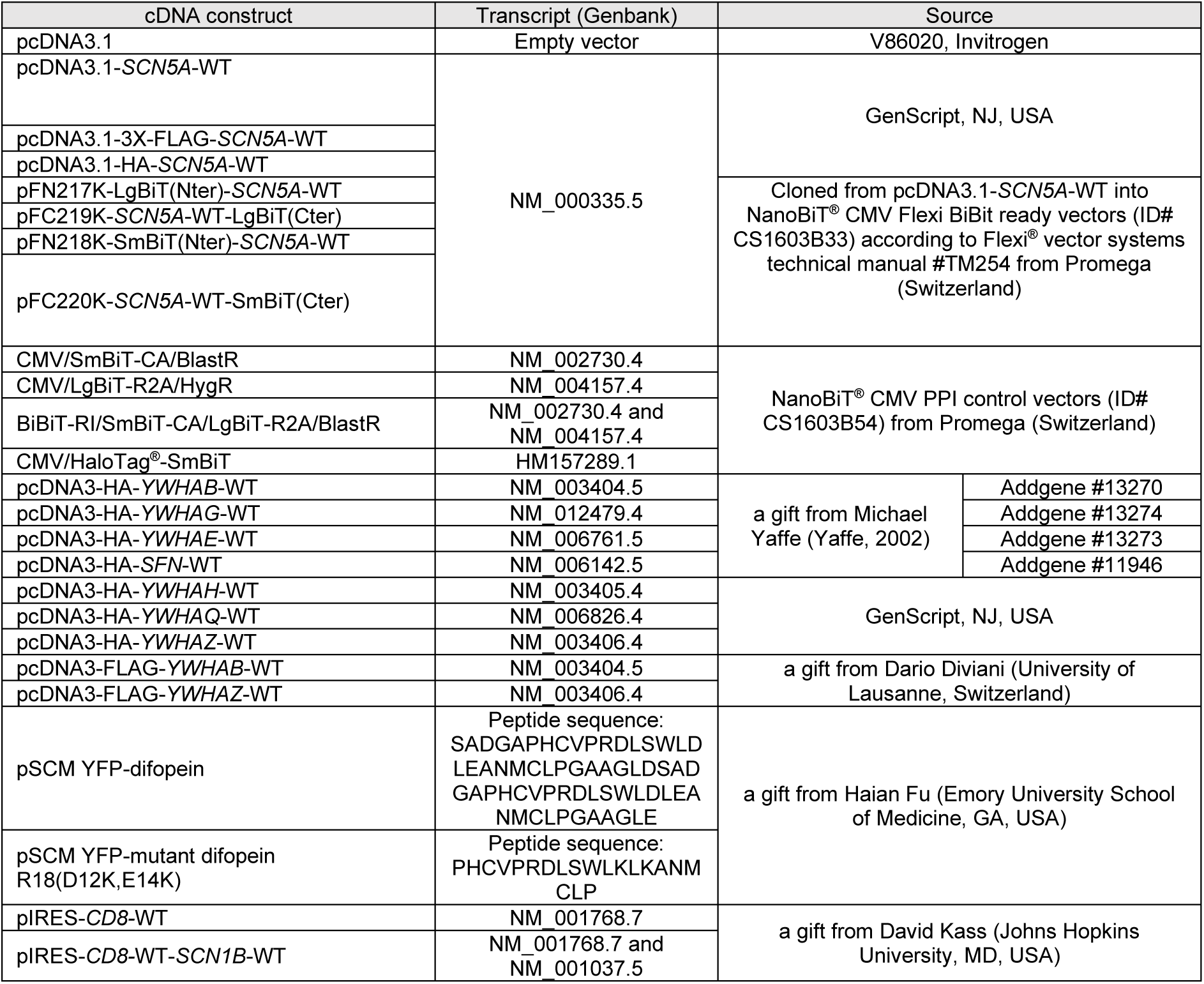
Description of cDNA constructs used in this study.

To clone *SCN5A*-WT into NanoBiT^®^ CMV Flexi BiBit ready vectors, SgfI and PmeI restriction sites were introduced, for which the primers were designed by the Flexi® Vector Primer Design Tool (https://ch.promega.com/techserv/tools/flexivectortool/) and purchased from Eurofins Genomics (Germany) (forward primer 5’-3’: GTC GGC GAT CGC CAT GGC AAA CTT CCT ATT ACC TCG G; reverse primer 5’-3’: GCG AGT TTA AAC CAC GAT GGA CTC ACG GTC CC).

### Cell culture and transfection

Human embryonic kidney cells tsA201 (ECACC Cat# 96121229, RRID:CVCL_2737) were cultured up to 20 passages at 37°C with 5% CO_2_ in Dulbecco’s modified Eagle’s culture medium (41965, Gibco^TM^, Thermo Fisher Scientific, USA), supplemented with 2 mM L-glutamine (G7513, Sigma), 50 U/mL penicillin-streptomycin (15140122, Gibco^TM^) and 10% heat-inactivated fetal bovine serum (10270-106, Lot: 2440045, Gibco^TM^). When needed, cells were split using phosphate-buffered saline (PBS, 10010-015, Gibco^TM^) and 0.05% Trypsin-EDTA (25300-054, Gibco^TM^). Mycoplasma contamination status was tested weekly with PCR Mycoplasma Test Kit I/C (PK-CA91-1096, Promokine, PromoCell GmbH, Germany). cDNA constructs were introduced into cells by transfection with LipoD293^TM^ (SL100668, SignaGen^®^ Laboratories). Cells were harvested 48 hours after transfection unless stated otherwise. Stable cell lines tsA201-*SCN5A*-WT and tsA201-FLAG-*SCN5A*-WT were established by polyclonal selection and were maintained in culture growth media with 200 µg/mL Zeocin™ Selection Reagent (R25005, Gibco^TM^).

### RNA extraction, reverse transcription polymerase chain reaction, and agarose gel electrophoresis

RNA extraction was performed on PBS-prewashed cell pellets using ReliaPrep^TM^ RNA cell Miniprep Systems according to the manufacturer’s instruction (Z6010, Promega). The concentration of extracted RNA was determined by NanoDrop™ One/One^C^ Microvolume UV-Vis Spectrophotometer (ND-ONE-W, Thermo Fischer Scientific™). Reverse transcription (RT) was performed using a High-Capacity cDNA Reverse Transcription kit (4375222, Applied Biosystems) followed by polymerase chain reaction (PCR) with GoTaq^®^ G2 Master Mix (M7822, Promega) on Biometra TRIO Touch Thermocycler (207072X, LabGene Scientific, Switzerland). The PCR protocol was as follows: initial denaturation at 95°C for 2 minutes, 35 cycles of denaturation at 95°C for 1 minute, annealing at 60°C for 45 seconds and elongation at 72°C for 45 seconds per kb, and a final extension of 72°C for 5 minutes. PCR products were stored at 4°C. Primers with their expected amplicon sizes are listed in Table 2.

**Table 2.**
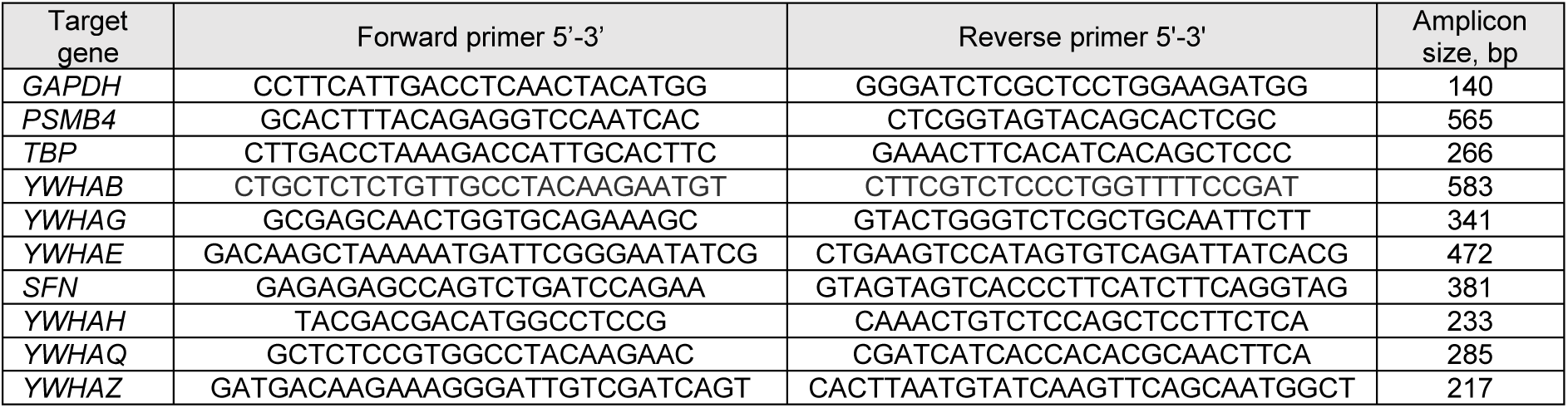
Description of primers used in this study.

### Protein lysate extraction

Pre-washed (PBS) adherent monolayers of cells were scraped in 2 mL of cold PBS (pH 7,4) and pelleted by centrifugation for 5 minutes at 200 g at 4°C. Cell pellets were lysed in lysis buffer (50 mM NaCl, 50 mM imidazole/HCl, 2 mM 6-aminohexanoic acid, 1 mM EDTA, pH 7) with addition of cOmplete tablets EDTA-free (04693132001, Roche, Switzerland), 0.5 mM Na_3_VO_4_, 0.5 mM NaFl, 10 μg/mL aprotinin, 10 μg/mL leupeptin, 1 mM phenylmethylsulfonyl fluoride and 1% digitonin (D141, Sigma) for 1 hour at 4°C. Lysates were centrifuged at 16,100 g for 15 minutes at 4°C. The obtained supernatant was taken for further analysis. Protein concentration was determined by Coo Assay Protein Dosage Reagent (Uptima, UPF86420).

### Co-immunoprecipitation

For immunoprecipitation nProtein A Sepharose 4 Fast Flow (17528001, Cytiva) or anti-FLAG^®^ M2 Magnetic Beads (M8823, Sigma) were used in the proportion of 1 μL of beads per 20 μg of total protein lysate. Protein lysates were diluted in Tris-buffered saline (TBS, 20 M Tris, 0.1 mM NaCl, pH 7.6, HCl-adjusted) and were mixed with the TBS-prewashed antibody-coupled beads at 4°C overnight. After thoroughly washing the beads with TBS, the co-immunoprecipitated protein complex was eluted with 4X LDS sample buffer (NP0007, Invitrogen). The samples were analyzed with the immunoblotting technique (see below).

### Cell surface biotinylation assay

Adherent cell monolayers were gently washed three times with cold PBS (pH 8) and incubated, while slowly rocking, with 1 mg/mL EZlink™ Sulfo-NHS-SS-Biotin (21331, Thermo Fisher Scientific) in PBS (pH 8) for 45 minutes at 4°C. Afterward, the excess biotin was quenched by washing with 50 mM Tris-HCl (pH 8), followed by several washes with PBS (pH 7,4). Extracted protein lysates were mixed with Streptavidin Sepharose High-Performance beads (GE Healthcare, USA) at 4°C overnight. After thoroughly washing the beads with washing buffer (0.1% BSA, 0.001% Tween20, PBS pH 7,4), the biotinylated fraction was eluted with 4X LDS sample buffer and analyzed with the immunoblotting technique (see below).

### Sodium dodecyl sulfate–polyacrylamide gel electrophoresis (SDS-PAGE) and immunoblotting analysis

Protein lysates containing LDS sample buffer and 100 mM dithiothreitol (DTT) were loaded onto 4-12% Tris-acetate acrylamide gels and let to run at 60 mV in Tris-acetate running buffer (50 mM Tris, 50 mM Tricine, 0.1% SDS, pH 8.3). Afterward, proteins were transferred to the nitrocellulose membrane by using the Trans-Blot Turbo system (1704158, Bio-Rad, USA). After validation of successful protein transfer with Ponceau S solution (0.1% Ponceau S, 12.5% acetic acid), the membranes were blocked with 5% bovine serum albumin (BSA) in TBS-T (TBS, 0.1% Tween20) for 1 hour at room temperature and then incubated with primary antibody dilutions at 4°C overnight. After incubating membranes with secondary antibodies for 1 hour at room temperature, the infrared fluorescent signals were revealed with LiCor Odyssey Infrared imaging system (LI-COR Biosciences) and quantified with ImageJ software (Rasband, W.S., U. S. National Institutes of Health, Bethesda, Maryland, USA).

Primary and secondary antibodies used in this study are listed in Table 3.

**Table 3.**
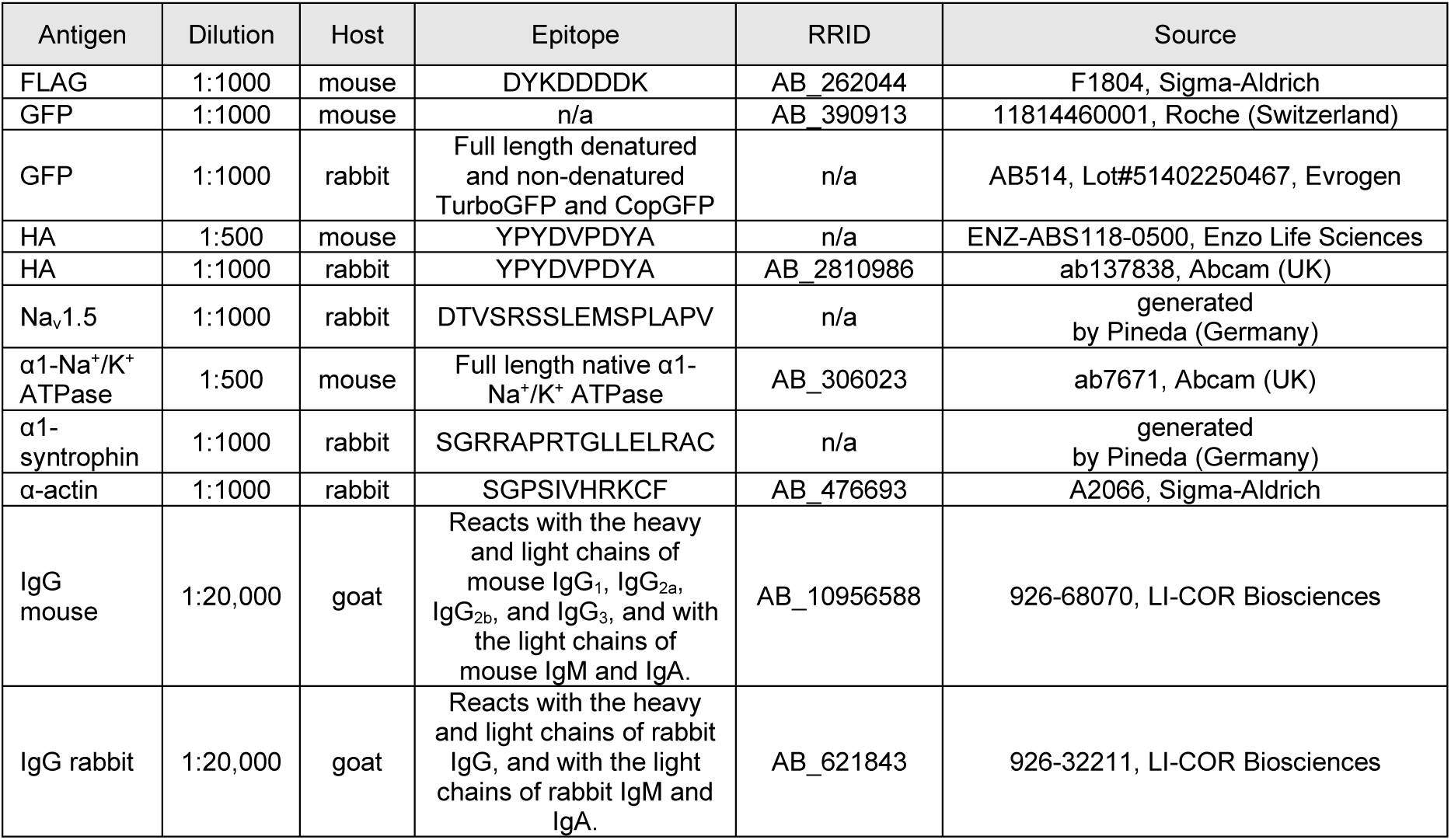
Description of primary and secondary antibodies used in this study.

### Patch clamp electrophysiology

To test data replicability, whole-cell sodium current recordings from two researchers (one of whom was blinded) using different setups were pooled together: either with an Axopatch 200B amplifier (Molecular Devices, Wokingham, United Kingdom) or with a VE-2 amplifier (Alembic Instrument, USA). Cells, solutions, micropipettes, and protocols were identical.

All cells were transiently co-transfected with *CD8*. Only successfully transfected cells, visualized by brief pre-wash with 1:1000 dilution of Dynabeads^TM^CD8 (Thermo Fisher Scientific), were chosen for electrophysiological recordings. Thin-wall capillaries (TW150F-3, World Precision Instruments, Germany) were pulled with a DMZ-universal puller (Zeitz, Germany) with a tip resistance of 1.0 to 3.0 MΩ. The intracellular pipette solution contained (in mM): CsAsp 70, CsCl 60, EGTA 11, CaCl_2_ 1, MgCl_2_ 1, HEPES 10, and 5 Na_2_ATP. The extracellular solution contained (in mM): NaCl 20, NMDG-Cl 110, CaCl_2_ 2, MgCl_2_ 1.2, HEPES 10, CsCl 5, D-glucose 5. pH was adjusted to 7.2-7.4 with CsOH or HCl. Osmolarity was measured by Osmometer type OM 806 (Löser) in mOsmol/kg H_2_O range. All recordings were performed at a stable temperature of 25°C using Axon™ pCLAMP™ 10 Electrophysiology Data Acquisition & Analysis Software, Version 10.2 (Axon Instruments, CA, USA). The recorded sodium current *I*_Na_ was filtered with a low-pass Bessel 5 kHz filter at a sampling rate of 20 kHz per signal. Recordings without leak subtraction were selected for analysis. Raw traces were not corrected for the liquid junction potential.

Exclusion criteria for individual traces were: absence of *I*_Na_; current at holding potential (-100 mV) <= -100 pA; slope of the activation (A) curve <=4; and slope of the steady-state inactivation (SSI) curve >=8. Sodium current density (pA/pF) was calculated by dividing peak current by cell capacitance. Current-voltage (I–V) curves were fitted with the equation *y* = *g*(*V*_m_ – *V*_rev_)/((1 + exp[(*V*_m_ – *V*_1/2_)/*k*])). SSI and A curves were fitted with the Boltzmann equation *y* = 1/(1 + exp[(*V*_m_ – *V*_1/2_)/*k*]), where *y* is the normalized current or conductance at a given holding potential, *V*_m_ is the membrane potential, *V*_rev_ is the reversal potential, *V*_1/2_ is the potential at which half of the channels are activated, and *k* is the slope factor. Data analysis was not blinded.

### Protein-protein interaction assay

The NanoLuc^®^ Binary Technology, NanoBiT^®^ (Promega) allowed us to monitor protein-protein interactions between two Na_v_1.5-WT subunits in live cells using an aqueous, cell-permeable furimazine substrate. For this purpose, *SCN5A*-WT was cloned into different vectors that encoded Na_v_1.5-WT tagged with Large BiT (LgBiT) and Small BiT (SmBiT) on N- or C-termini when transfected into tsA201 cells. In the case of physical interaction between LgBiT and SmBiT, the bright luminescent signal was formed. Since Nano-Glo^®^ Live Cell Assay (N2012, Promega) was not ratiometric and depended on the cell quantity, we normalized the luminescent signal to the fluorescent signal obtained with CellTiter-Fluor^®^ Cell Viability assay (G6082, Promega). The signals were detected with Spark^®^ multimode microplate (Tecan) and GloMax^®^ Explorer Multimode Microplate Reader (GM3500, Promega).

### Data and statistical analysis

Data are represented as means ± SEM. Data normality was tested by using Shapiro-Wilk test. Statistical significance for normally distributed data was calculated with ordinary one-way ANOVA. In case data were normalized to control condition statistical significance was calculated with one-sample two-tailed t-test with hypothetical mean value=1. Statistical significance for non-normally distributed data was calculated with Kruskal-Wallis test (Prism version 7.04; GraphPad, CA, USA). Exact *p*-values are indicated in the figures.

### Supplemental material

Suppl. fig. S1-S2 show blots for co-immunoprecipitation between each of 14-3-3 isoforms and Na_v_1.5 in tsA201 cell line. Suppl. fig. S3 shows endogenous expression of all 14-3-3 genes in tsA201 cell line. Suppl. fig. S4 shows validation of difopein as an efficient competitor for 14-3-3/ligand interaction in tsA201 cells: difopein, but not the mutant of difopein, interacts with 14-3-3 and stabilizes its oligomerization.

## Results

### Identification of 14-3-3 isoforms interacting with Na_v_1.5

To identify which 14-3-3 isoform interacts with Na_v_1.5, we performed co-immunoprecipitation analysis. Individual overexpression of all seven human 14-3-3 isoforms tagged with HA revealed that by themselves 14-3-3 proteins have a certain degree of non-specific binding with the anti-FLAG magnetic beads (Suppl. fig. S1-2). Accordingly, the control condition with non-tagged Na_v_1.5 was run along with FLAG-Na_v_1.5 and compared for the intensity of co-immunoprecipitated 14-3-3 protein. As an indicator of successfully co-immunoprecipitated Na_v_1-5-specific fraction, we used α1-syntrophin, a known partner of Na_v_1.5 macromolecular complex and endogenously present in tsA201. From all the isoforms, only 14-3-3η exhibited reproducibly brighter band intensity in the FLAG-immunoprecipitated fraction compared to the non-tagged fraction; hence, we concluded that only this isoform reliably interacts with Na_v_1.5 (Suppl. fig. S1-2). Furthermore, sepharose beads coupled with anti-Na_v_1.5 antibody successfully co-immunoprecipitated α1-syntrophin and 14-3-3η, but not 14-3-3σ (Fig. 1). However, due to the overall weakness of Na_v_1.5-immunoprecipitated 14-3-3η protein, we suggest that the interaction is rather weak and/or transient.

**Figure 1.**
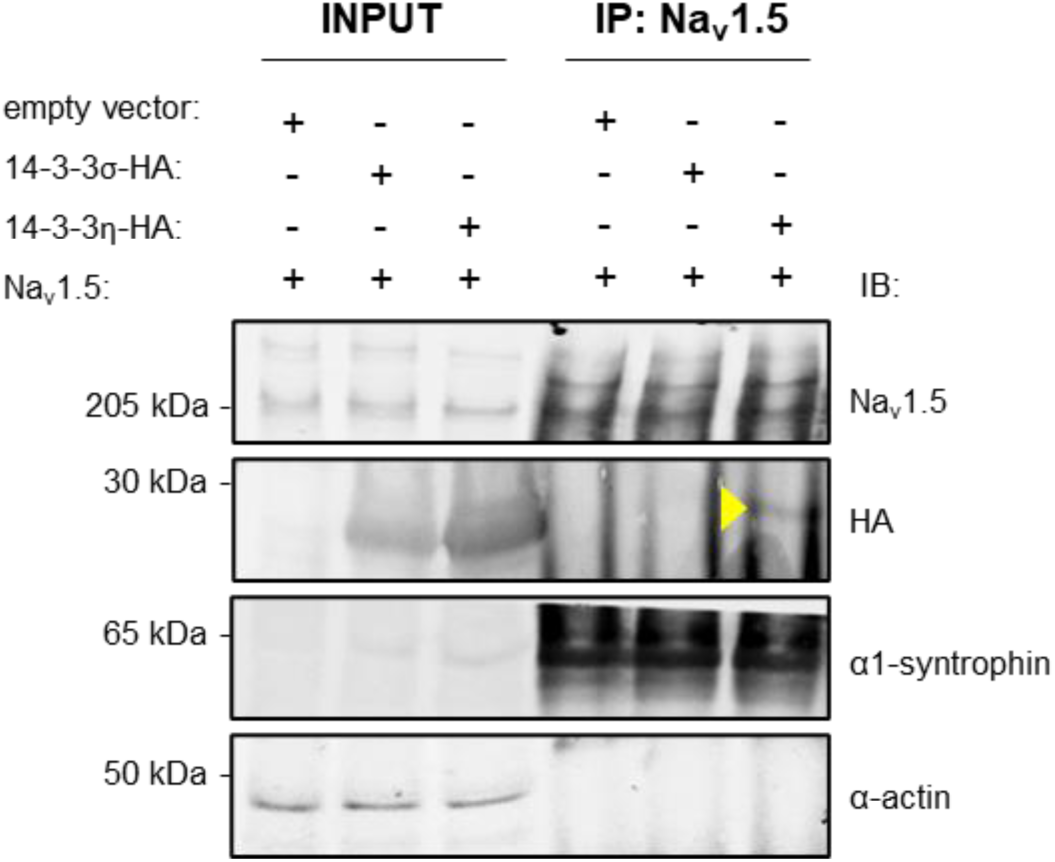
Na_v_1.5 co-immunoprecipitated with 14-3-3η. Immunoblot of total lysate and Na_v_1.5-specific immunoprecipitated fraction of tsA201 cells 48 hours after transient co-expression of an empty vector, 14-3-3σ, 14-3-3η and Na_v_1.5. Immunoprecipitation was performed with Sepharose beads pre-coupled with anti-Na_v_1.5 antibody. HA-tagged 14-3-3σ and 14-3-3η were revealed with an anti-HA antibody. Endogenous α1-syntrophin was used as a positive control for co-immunoprecipitation with Na_v_1.5, and α-actin as a negative control. Yellow triangle indicates on the band corresponding to the co-immunoprecipitated 14-3-3η.

### 14-3-3 proteins do not affect expression level of Na_v_1.5 at the cell surface

Next, we assessed the role of 14-3-3 proteins on the cell surface density of Na_v_1.5. This was motivated by findings from Pohl *et al*., who suggested that 14-3-3 proteins modulate cell surface expression of membrane receptors and ion channels upon binding with the E3 ubiquitin ligase Nedd4-2 (Pohl et al., 2021). Specifically, they showed that the high affinity of 14-3-3 with phosphorylated Ser342 and Ser448 sites of Nedd4-2 induces its structural rearrangement with plausible exposure of its tryptophan-rich WW-domain (Pohl et al., 2021). Our group previously reported the interaction between the WW-domain of Nedd4-2 with the PY-motif of Na_v_1.5 and its importance for Na_v_1.5 cell surface localization (Van Bemmelen et al., 2004; Rougier et al., 2005). According to our current results, the levels of the total and cell surface Na_v_1.5 expression, represented by the biotinylated fraction, did not change upon overexpression of each individual 14-3-3 isoform when compared to the sodium channel alone (Fig. 2A-B).

**Figure 2.**
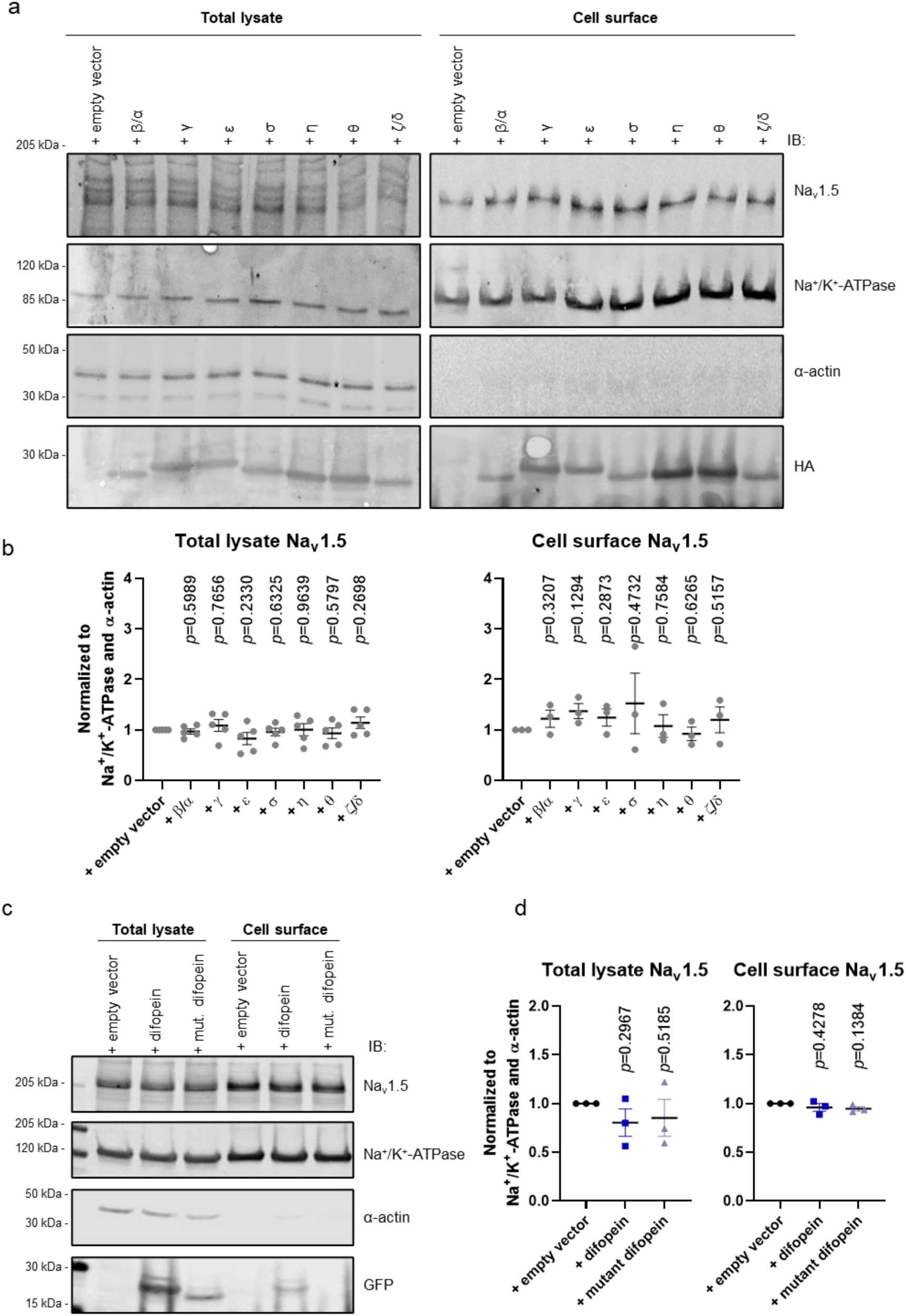
14-3-3 proteins do not affect the total expression level of Na_v_1.5 and cell surface density. Representative immunoblots of the total lysate and cell surface fraction of tsA201 stably expressing Na_v_1.5 after 48 hours of transient overexpression of: **(a)** HA-tagged 14-3-3 proteins; **(c)** YFP-tagged difopein and its mutant. Endogenous Na^+^/K^+^-ATPase was used as a positive control of cell surface fraction, while α-actin was used as a negative control. **(b** and **d)** Intensities of Na_v_1.5 in total protein lysate and at the cell surface were normalized to Na^+^/K^+^-ATPase and α-actin and to the control condition (“+ empty vector”). Data are presented as mean ± SEM from three-five biological replicates. Individual *p*-values, calculated with one-sample two-tailed t-test with hypothetical mean value=1, are indicated in each panel.

Of note, in tsA201, we detected endogenous mRNAs of all 14-3-3 isoforms (Suppl. fig. S3). In line with our results, transcripts of all 14-3-3 isoforms were previously reported for HEK293, the cell line from which tsA201 was originally derived (Mathew et al., 2016). Therefore, to inhibit endogenous 14-3-3/ligand interactions, we used difopein, a dimer of R18, the unphosphorylated peptide antagonist of 14-3-3 (Masters and Fu, 2001). Difopein strongly interacts with two 14-3-3 proteins, stabilizing them in dimeric conformation, occupying their binding grooves, and preventing their interaction with other ligands (Masters and Fu, 2001). We performed co-immunoprecipitation analysis to validate that difopein is an effective competitor for 14-3-3/ligand interactions in our heterologous expression system. First, we confirmed that 14-3-3 strongly interacts with difopein but not with its mutant (Suppl. fig. S4a). As expected, difopein-YFP exhibited higher molecular weight than the mutant difopein-YFP, represented by the monomeric R18-D12K,E14K peptide (Suppl. fig. S4a). Further, we demonstrated that 14-3-3 dimers were stabilized by difopein but not by its mutant (Suppl. fig. S4b). Last, we confirmed that such an effect did not depend on any specific 14-3-3 isoform (Suppl. fig. S4b). Of note, difopein and its mutant did not affect the total and cell surface expression of Na_v_1.5 (Fig. 2c-d).

Therefore, we concluded that 14-3-3 proteins do not regulate Na_v_1.5 expression and cell surface density in the tested heterologous expression system.

### Impact of 14-3-3 proteins onto Na_v_1.5-mediated sodium current

Since of all 14-3-3 isoforms, only 14-3-3η weakly but reproducibly interacted with Na_v_1.5 (Fig. 1, Suppl. fig. S1-S2), we assessed the whole cell recordings of the sodium current (*I_Na_*) when transiently overexpressing *YWHAH*, encoding 14-3-3η. Moreover, to establish comparable conditions with the previous study that described a 14-3-3η-induced negative shift in *I_Na_* inactivation curve, we similarly co-expressed *SCN1B*, encoding the Na_v_β1 subunit of the voltage-gated sodium channel (Allouis et al., 2006). In our heterologous expression system, neither 14-3-3η or Na_v_β1 alone nor their combination modified the baseline whole-cell properties of *I_Na_*, as described by the current-voltage (*I*-*V*) relationship, peak current density, reversal potential (*V*_rev_) and half-maximal potentials (*V*_1/2_) of activation and inactivation curves (Fig. 3, Table 4). Similarly, disruption of endogenous 14-3-3/ligand interactions with difopein did not modify basal current properties of Na_v_1.5 (Fig. 4, Table 5).

**Figure 3.**
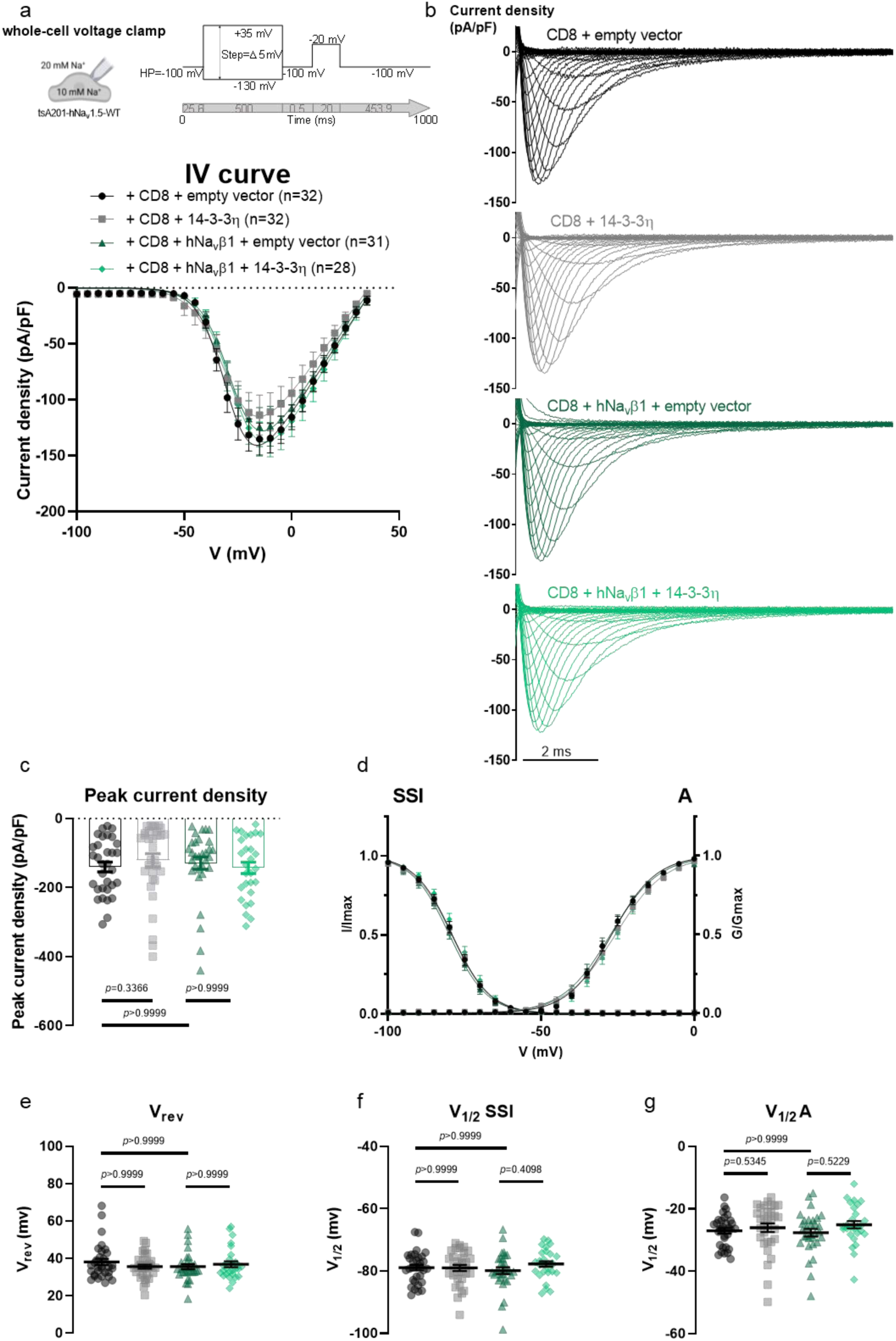
14-3-3η expressed alone or with Na_v_β1 does not affect Na_v_1.5-conducted sodium current. **(a)** Current density-voltage (IV) relationships from tsA201-Na_v_1.5 transiently expressing CD8 with empty vector, or 14-3-3η, and/or Na_v_β1. **(b)** Representative whole-cell *I*_Na_ traces recorded from the listed conditions. **(c)** Peak current densities of each group. **(d)** Activation (A) and steady-state inactivation (SSI) curves. **(e)** Reversal potential (*V*_rev_). **(f)** Half-voltage of SSI (*V*_1/2_ SSI). **(g)** Half-voltage of A (*V*_1/2_ A). Values pertaining to the biophysical properties are shown in Table 4. Data are presented as mean ± SEM. *n*, number of cells taken for analysis. Kruskal-Wallis has been performed in **(c, e, f,** and **g)** and the multiplicity adjusted *p*-values are indicated in each panel.

**Figure 4.**
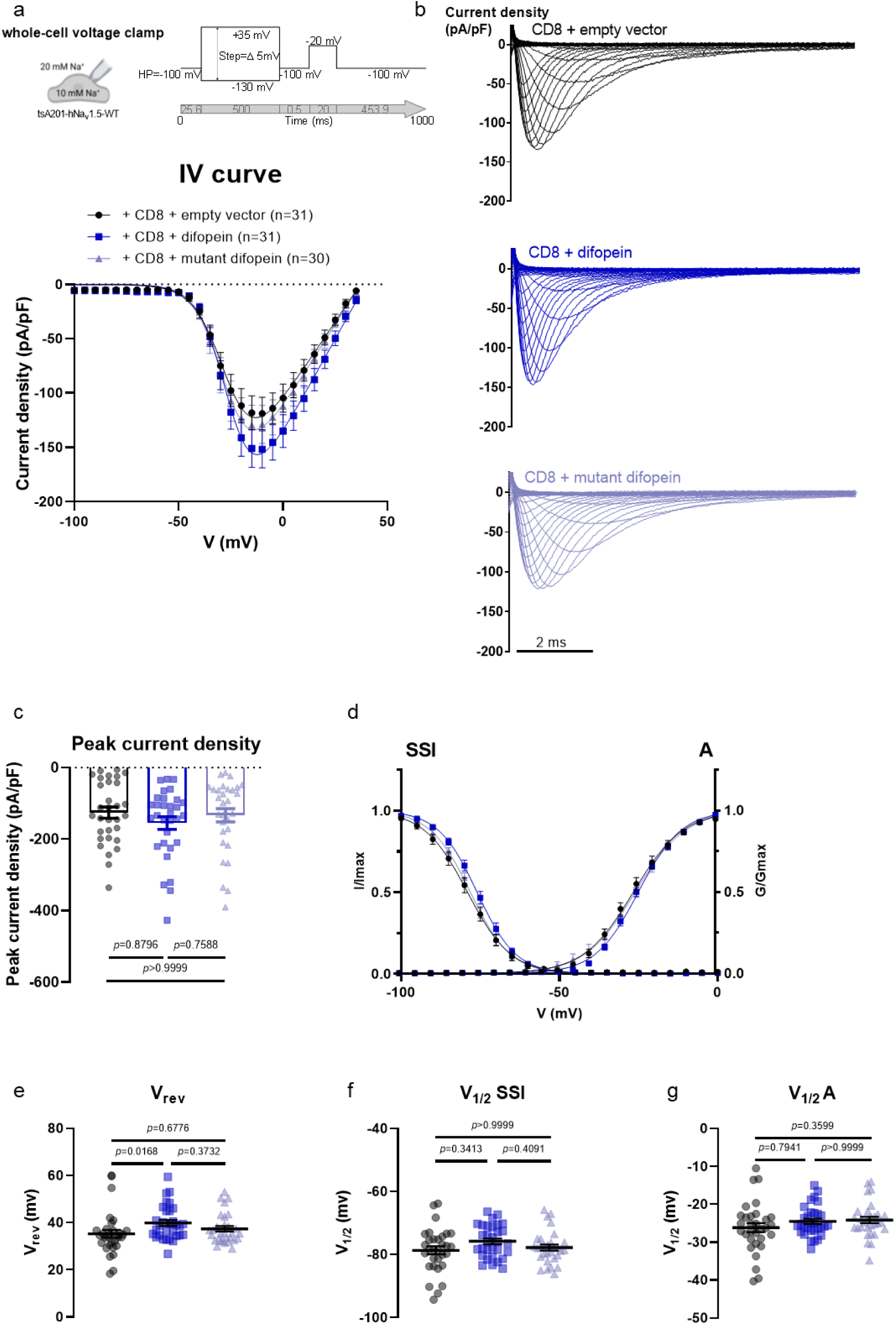
Difopein, an antagonist of 14-3-3/ligand interactions, does not affect Na_v_1.5-conducted sodium current. **(a)** Current density-voltage (IV) relationships from Na_v_1.5-expressing tsA201 cells transiently expressing CD8 with empty vector, difopein, or mutant of difopein. **(b)** Representative whole-cell *I*_Na_ traces recorded from the listed conditions. **(c)** Peak current densities of each group. **(d)** Activation (A) and steady-state inactivation (SSI) curves. **(e)** Reversal potential (*V*_rev_). **(f)** Half-voltage of SSI (*V*_1/2_ SSI). **(g)** Half-voltage of A (*V*_1/2_ A). Values pertaining to the biophysical properties are shown in Table 5. Data are presented as mean ± SEM. *n*, number of cells taken for analysis. Kruskal-Wallis has been performed in **(c, e, f,** and **g)** and the multiplicity adjusted *p*-values are indicated in each panel.

**Table 4.** Biophysical properties of sodium currents conducted by Na_v_1.5 stably present in tsA201 cells in combination with transiently expressed 14-3-3η and/or Na_v_β1. CD8 was co-expressed to visualize successfully transfected cells. Peak sodium current (I_Na_) densities are extracted from the respective I-V curves. Data are presented as mean ± SEM and are compared to control values (i.e., “+CD8 + empty vector”). n, number of cells analyzed.

| | Peak $I_{Na}$ density<br>(pA/pF) | $V_{rev}$ (mV) | Steady-state activation | | Steady-state inactivation | |
| --- | --- | --- | --- | --- | --- | --- |
| | | | $k$ slope | $V_{1/2}$ (mV) | $k$ slope | $V_{1/2}$ (mV) |
| + CD8<br>+ empty vector<br>(n=32) | -140.18 ± 14.11 | 38.15 ± 1.68 | 6.38 ± 0.17 | -27.01 ± 0.89 | 5.25 ± 0.12 | -78.93 ± 0.90 |
| + CD8 + 14-3-3η<br>(n=32) | -120.69 ± 19.06 | 35.62 ± 1.13 | 6.45 ± 0.18 | -26.02 ± 1.38 | 5.22 ± 0.16 | -78.98 ± 0.97 |
| + CD8 + h$Na_v\beta1$<br>+ empty vector<br>(n=31) | -129.51 ± 17.83 | 35.59 ± 1.35 | 6.49 ± 0.13 | -27.61 ± 1.22 | 4.97 ± 0.14 | -79.86 ± 1.10 |
| + CD8 + h$Na_v\beta1$<br>+ 14-3-3η<br>(n=28) | -143.15 ± 16.60 | 36.88 ± 1.57 | 6.39 ± 0.17 | -25.05 ± 1.18 | 5.35 ± 0.17 | -77.75 ± 0.90 |

**Table 5.**
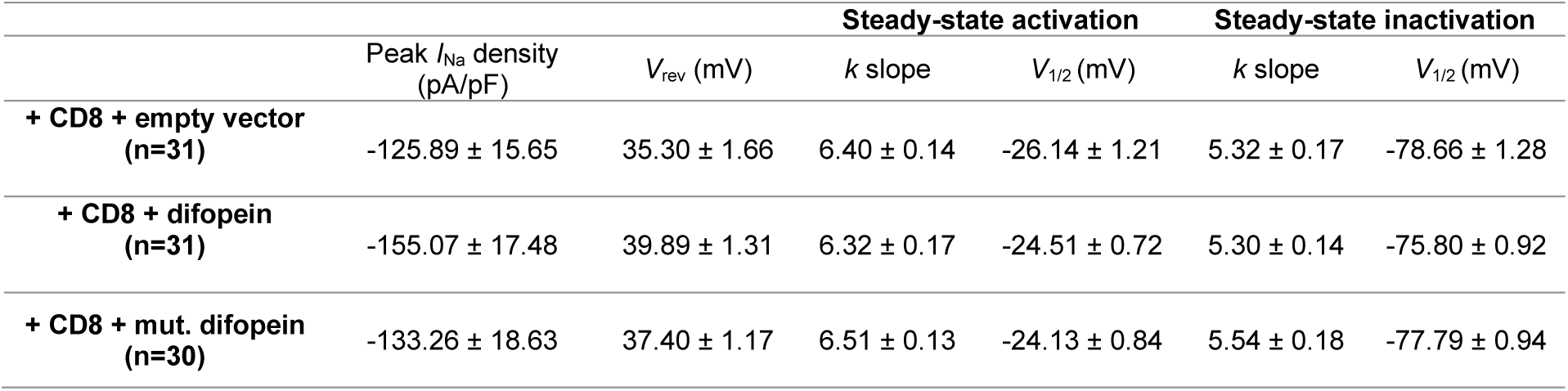
Biophysical properties of sodium currents conducted by Na_v_1.5 stably present in tsA201 cells in combination with transiently expressed difopein and its mutant. CD8 was co-expressed to visualize successfully transfected cells. Peak sodium current (I_Na_) densities are extracted from the respective I-V curves. Data are presented as mean ± SEM and are compared to control values (i.e., “+CD8 + empty vector”). n, number of cells analyzed.

In summary, the isoform non-specific antagonist of 14-3-3, difopein, and 14-3-3η, the only isoform interacting with Na_v_1.5, left the channel function unmodified.

### Inhibition of 14-3-3 dimerization does not affect the interaction between α-subunits of Na_v_1.5

One role of the 14-3-3 family of adapter proteins is to stabilize multiprotein complexes (Obsil and Obsilova, 2011). In particular, 14-3-3η was proposed to mediate the coupled gating of Na_v_1.5 dimers by bridging two α-subunits through their DI-DII loops (Clatot et al., 2017). We evaluated whether the 14-3-3 antagonist, difopein, affected Na_v_1.5 dimerization. Our co-immunoprecipitation analysis confirmed the existence of Na_v_1.5-Na_v_1.5 interactions in a heterologous expression system, but these were not affected by difopein overexpression (Fig. 5a-b). To confirm this finding in a more physiological and dynamic environment, we performed a NanoBiT assay. In brief, N- and C-termini of Na_v_1.5 were tagged with LgBiT and SmBiT, respectively, which are parts of the bright and stable Nanoluciferase. Accordingly, when two Na_v_1.5 physically interact, LgBiT and SmBiT would reconstitute Nanoluciferase, subsequently leading to luminescence emission (Fig. 5c). Even though intrinsic interaction between LgBiT and SmBiT is rather weak (*K*_d_ = 190 μM) (Dixon et al., 2016), all experiments were accompanied by a simultaneous co-expression of Na_v_1.5-LgBiT with the non-interacting HaloTag-SmBiT as a readout of the background luminescence (Fig. 5c). In live tsA201 cells, the level of interaction between two Na_v_1.5 was not modified by difopein (Fig. 5d-g). Interestingly, when comparing various locations of LgBiT and SmBiT tags on Na_v_1.5, the brightest luminescent signal was detected when N-N and C-C termini were combined, compared to the N-C and C-N conditions (Fig. 5d-g). It might indicate a preferred conformational orientation of α-subunits within Na_v_1.5 dimers.

**Figure 5.**
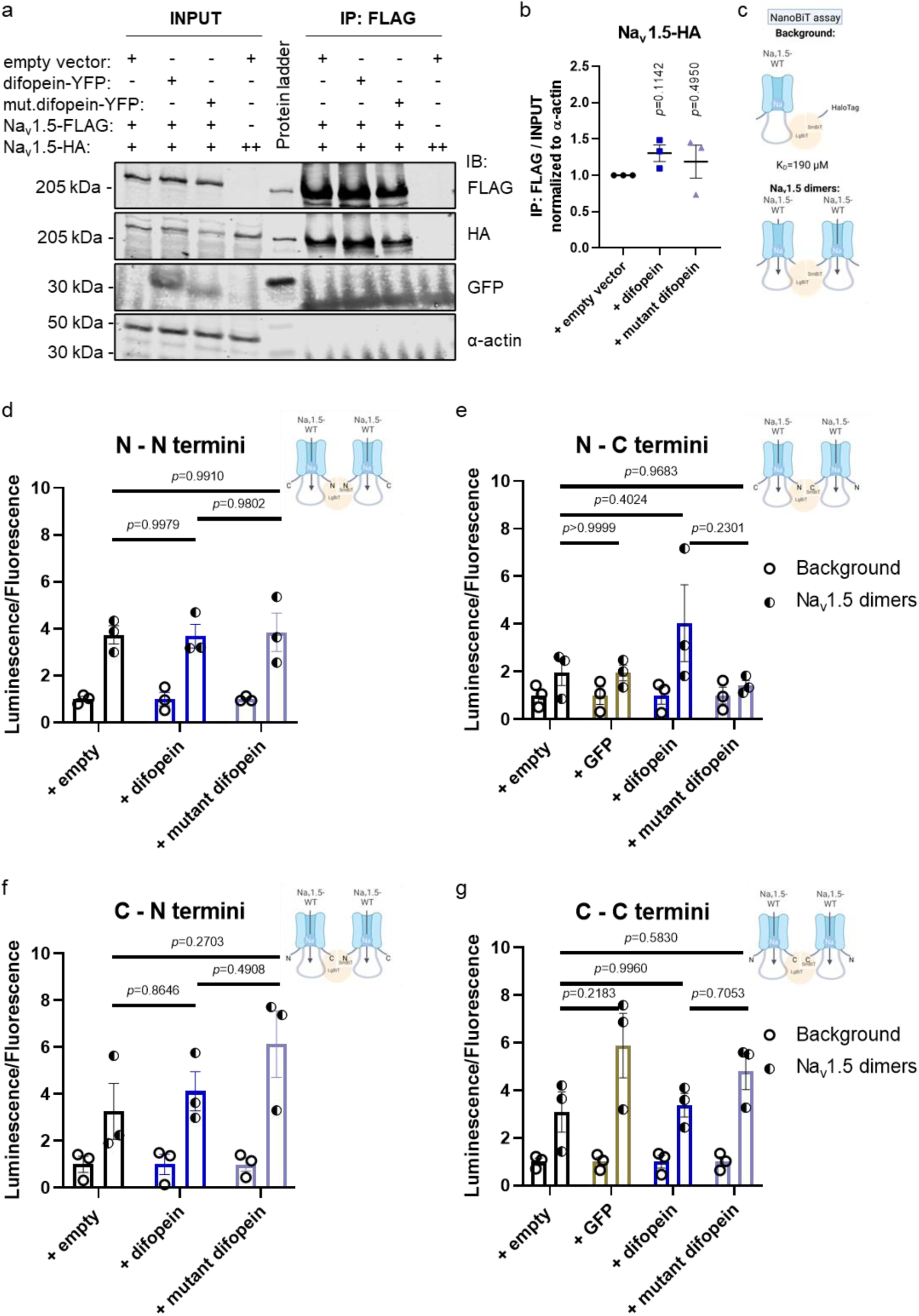
Inhibition of 14-3-3/ligand interaction does not affect oligomerization between α-subunits of Na_v_1.5. Difopein and its mutant did not affect co-immunoprecipitation between α subunits of Na_v_1.5. **(a)** Representative immunoblot of the total lysate (“INPUT”) and FLAG-specific immunoprecipitated fraction (“IP: FLAG”) of tsA201 cells transiently expressing Na_v_1.5 with empty vector, difopein, or mutant of difopein. Overexpressed YFP-tagged difopein and its mutant were revealed with an anti-GFP antibody. Endogenous α-actin was used as a negative control for co-immunoprecipitation with Na_v_1.5. **(b)** Intensity of co-immunoprecipitated Na_v_1.5-HA was normalized to the intensity of the total Na_v_1.5-HA divided by the intensity of α-actin. Data are presented as mean ± SEM from three biological replicates and are normalized to the control condition (“+ empty vector”). Individual *p*-values, calculated with one-sample two-tailed t-test with hypothetical mean value=1, are indicated in the panel. **(c)** Schematic illustration of NanoBiT assay. Background signal was determined by co-expression of N- or C-termini LgBiT-Na_v_1.5 with non-interacting control, SmBiT-HaloTag. **(d, e, f,** and **g)** In living cells difopein did not modify the level of Na_v_1.5 dimers. Results of the NanoBiT assay are presented as relative intensity of luminescence (indicating the level of protein-protein interactions) normalized to fluorescence (indicating the number of cells) in tsA201 cells transiently transfected with *SCN5A* constructs. Each dataset was normalized to its relevant background. Different variations between Na_v_1.5 N- and C-termini interactions were tested. Since difopein and its mutant were tagged with YFP, we also included control by co-transfection with GFP. NT, non-transfected. Data are presented as mean ± SEM from three biological replicates. The multiplicity adjusted *p*-values, calculated with one-way ANOVA, are indicated in each panel.

## Discussion

We reported that amongst seven 14-3-3 isoforms, only η (encoded by *YWHAH* gene) weakly co-immunoprecipitated with the major cardiac voltage-gated sodium channel Na_v_1.5 when heterologously expressed in tsA201 cells. Nevertheless, neither 14-3-3η overexpression nor its inhibition with difopein affected the total protein level of Na_v_1.5, its cell surface localization, dimerization, and basal biophysical properties of *I_Na_*.

Our finding that 14-3-3η interacted with Na_v_1.5 is in line with previous report of Allouis *et al*., who demonstrated that 14-3-3 interacted with Na_v_1.5 in Cos-7 and native mouse cardiac tissue and identified by yeast 2-hybrid screen DI-DII loop as a specific binding region of 14-3-3η (Allouis et al., 2006). Of note, when all cardiac voltage-gated sodium channels, including Na_v_1.5, were immunoprecipitated from the mouse left ventricles using pan-Na_v_ antibody, the most abundant peptides in the fraction were corresponding to 14-3-3ζ/δ, 14-3-3ε, 14-3-3γ, 14-3-3β/α, and 14-3-3σ, but not to 14-3-3η (Lorenzini et al., 2021).

Furthermore, we reported that difopein and 14-3-3η did not modify *I_Na_* density and gating kinetics. Likewise, *I_Na_* density was not modified by exogenous 14-3-3η, by inhibition of endogenous 14-3-3η, and by isoform-unspecific antagonist, difopein (Allouis et al., 2006; Utrilla et al., 2017; Zheng et al., 2021). However, regarding the activation and inactivation kinetics of Na_v_1.5, several discrepancies were described. Similarly with our study, 14-3-3η overexpression did not modify conductance-voltage relationship (Allouis et al., 2006). But in contrast with our study, Zheng *et al*. demonstrated that difopein induced negative shift of the voltage-dependent conductance curve for the wild-type Na_v_1.5 as well as for its truncated mutant Gly1642X with His558Arg polymorphism (Zheng et al., 2021). Furthermore, unlike our study, Allouis *et al*. showed that 14-3-3η shifted the inactivation curve to more negative potentials and decelerated the recovery from the inactivation of Na_v_1.5 (Allouis et al., 2006). By co-expressing *SCN1B*, we eliminated the possibility that such discrepancy could be due to the presence of the Na_v_β1 subunit. In fact, we did not observe any significant effects of *SCN1B* expression onto biophysical properties of the heterologous Na_v_1.5. Indeed, previous studies reported conflicting results regarding the impact of Na_v_β1 on the function of Na_v_1.5. While some groups detected changes of the peak current density and/or differences in the channel kinetics and gating properties, others like us did not observe any Na_v_β1-mediated changes of *I_Na_* (Malhotra et al., 2001; Maroni et al., 2019; Martinez-Moreno et al., 2020; Valdivia et al., 2002; Zhu et al., 2021). Unlike the tsA201 cells used in our study, Allouis *et al*. used Cos-7 cells derived from monkey kidney fibroblasts as their heterologous expression system (Allouis et al., 2006). Like tsA201 cells, Cos-7 endogenously express all seven 14-3-3 isoforms, too (Abdrabou et al., 2020). However, their expression pattern, representing subcellular localization of 14-3-3 proteins, differed noticeably from human embryonic kidney (HEK293) cells (Abdrabou et al., 2020). Furthermore, other protein partners that regulate the function of Na_v_1.5 may also vary between cellular models and may explain the disparity of our outcomes.

Our observation that 14-3-3 proteins do not participate in the dimerization of heterologously expressed Na_v_1.5 was also corroborated by previous reports (Clatot et al., 2017; Utrilla et al., 2017). Both studies highlighted that instead of directly mediating the formation of Na_v_1.5 homo- and heterocomplexes with other ion channels (*e.g.*, Na_v_1.5-Na_v_1.5 and Na_v_1.5-K_ir2.1_), 14-3-3η played a significant role in their biophysical coupling (Clatot et al., 2017; Utrilla et al., 2017).

Zheng *et al*. also suggested additional indirect roles for 14-3-3-dependent regulation of Na_v_1.5 (Zheng et al., 2021). Indeed, since Na_v_1.5 is regulated by many proteins, including kinases, of which many are 14-3-3 substrates, too (Abriel, 2010; Iqbal and Lemmens-Gruber, 2019), it is plausible that 14-3-3 proteins could alter the function of Na_v_1.5 indirectly through other partner proteins (Fig. 6). Indeed, monomeric peptide R18, of which difopein is composed, disrupts 14-3-3γ/ε from protein kinase A (PKA), unmasking PKA’s catalytic activity and promoting PKA-mediated phosphorylation of its downstream targets (Kent et al., 2010). Na_v_1.5 is among PKA’s targets, and PKA stimulation in HEK293 cells has been shown to enhance *I_Na_* (Aiba et al., 2014). Furthermore, 14-3-3γ was shown to slow dephosphorylation of calcium/calmodulin-dependent protein kinase (CaMKII) (Psenakova et al., 2018). CaMKII is reported to mediate both gain- and loss-of-function effects of *I_Na_*, highlighting its complex relationship with Na_v_1.5 (Iqbal and Lemmens-Gruber, 2019). Moreover, 14-3-3ζ has been shown to regulate protein kinase C (PKC), which, in turn, also modulates Na_v_1.5 activity through phosphorylation (Gannon-Murakami and Murakami, 2002). Nevertheless, in our heterologous expression system the effect of isoform-unspecific antagonist, difopein, on the regulation of Na_v_1.5 was not evident.

In summary, ubiquitous presence of homo- and hetero-oligomerizing 14-3-3 proteins makes investigation of their direct role onto Na_v_1.5 rather challenging. Furthermore, expression of 14-3-3 proteins and their downstream targets might drastically vary between different models, potentially providing for distinct direct and indirect modes of Na_v_1.5 regulation.

### Limitations

To eliminate possible clonal specificity of our cell model, we developed polyclonal tsA201 stably expressing Na_v_1.5. This led to high variability in whole-cell Na_v_1.5 expression, which made single-cell electrophysiological analyses challenging, we strengthened the reliability of our data by testing a relatively large number of cells for each condition.

One of the limitations of this study is the presence of all endogenous 14-3-3 isoforms, including 14-3-3η, that could saturate Na_v_1.5-binding sites and hence preclude interaction with the overexpressed proteins. In addition to homodimerization 14-3-3 isoforms are also capable to form heterodimers (Jones et al., 1995; Obsil and Obsilova, 2011). Thus, it is possible that even when overexpressed 14-3-3η-comprised heterodimers may interact with substrates other than Na_v_1.5, hindering its direct role on the channel. Therefore, using gene-editing techniques to silence specific isoforms would be advantageous to unravel the complexity of 14-3-3 signaling.

Another limitation of heterologously expressed proteins is the difficulty in controlling post-translational modifications, such as phosphorylation, that impact Na_v_1.5 function and 14-3-3 signaling. A recent study reported a successful semisynthetic approach to stabilize Na_v_1.5 phosphorylation *in vitro* (Galleano et al., 2021). Thus, it would be interesting to test 14-3-3-specific regulation under stably phosphorylated Na_v_1.5.

While 14-3-3 might not act directly onto Na_v_1.5, it was shown to act onto its macromolecular complex with other protein partners (*e.g.*, PKA, PKC, CaMKII) and cardiac ion channels (*e.g.*, Kir2.1) (Pohl et al., 2021; Utrilla et al., 2017). Because of the vast spectrum of 14-3-3 targets, one should be aware of the possible indirect effects measured in any experiment targeting 14-3-3 function. As such, any results obtained with difopein should be interpreted carefully.

In summary, while direct effects of 14-3-3, specifically isoform η, on the cardiac sodium channel are not evident in the tested heterologous expression system, it still may regulate Na_v_1.5 through proteins of its macromolecular complex.

## Conclusions

We confirmed that 14-3-3η forms a macromolecular complex with human Na_v_1.5 in a heterologous expression system. Difopein is a potent tool to antagonize 14-3-3-ligand interactions but not the direct physical interaction between two Na_v_1.5 α-subunits. Overexpression of 14-3-3η and difopein did not affect *I_Na_* density, gating kinetics as well as total and cell surface expression of Na_v_1.5. Overall, the role of 14-3-3 proteins on the functionality of Na_v_1.5 is dispensable in tsA201 cell line.

## Author contributions

**O.I**, **D.R.K**., **J.S.R.**, and **H.A.** designed this research project. **O.I.**, **A.F.H.**, **E.R.**, **A.S.**, **D.R.K.**, **N.S.**, **M.E.**, **S.G.**, and **J.S.R.** performed the experiments. **O.I.**, **A.F.H.**, and **J.S.R.** analyzed the data. **O.I.** and **A.F.H.** made figures and tables. **O.I.** drafted and edited the manuscript. **O.I.**, **J.S.R.**, and **H.A.** critically reviewed the manuscript.

## Disclosure statement

The authors declare no potential conflict of interest.

## Funding

This work was funded by the Swiss National Science Foundation [SNF 310030_184783] to H.A.

## Acknowledgments

We thank Dr Zoja Selimi (University of Bern) and Prof Jan Kucera (University of Bern) for their invaluable insights and fruitful discussions. We thank Dr Sarah Vermij for proofreading of the article.

## Notes

### Competing Interest Statement

The authors have declared no competing interest.

### Summary of Updates

We identified that the differences in Vrev between the two datasets of electrophysiological recordings were caused by one specific batch of the intracellular solution, used mainly in cells presented in Figure 4. Therefore, we excluded these data and performed additional electrophysiological recordings for the aforementioned conditions. In contrast to the previous version, difopein did not affect the biophysical properties of INa. We have also improved statistical analysis and representative immunoblots. Table 5, Figure 1, Figure 2, Figure 4, Figure 5 and Supplemental Figures were updated.

