## Supplemental Figures S1-S4 for "The dispensability of 14-3-3 proteins for the regulation of human cardiac sodium channel Na_v_1.5"

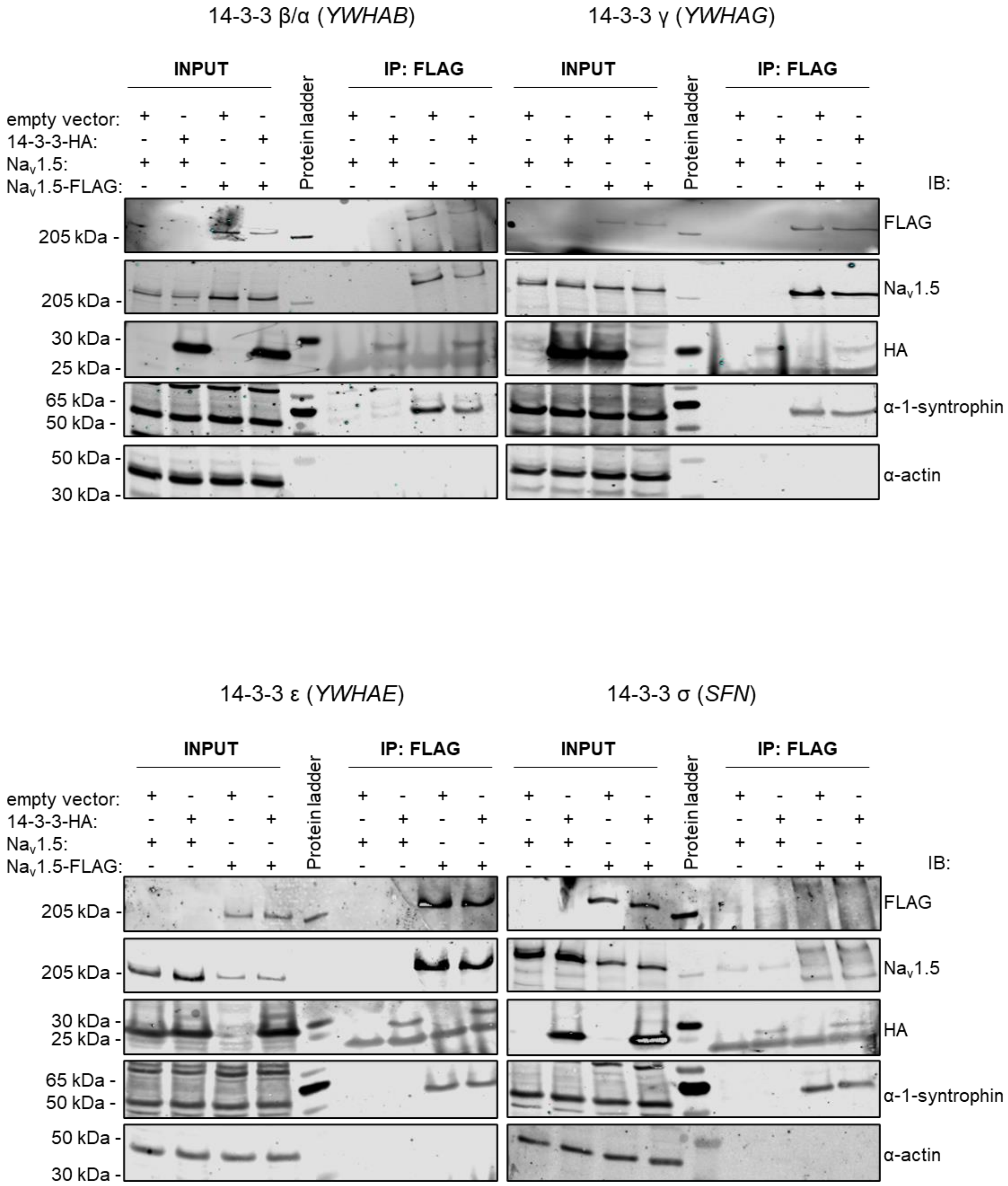


**Supplemental figure S1. Co-immunoprecipitation between $Na_V1.5$ and 14-3-3 isoforms:** 48 hours after transient overexpression of 14-3-3 β/α (encoded by *YWHAB*), 14-3-3 γ (encoded by *YWHAG*), 14-3-3 ε (encoded by *YWHAE*), 14-3-3 σ (encoded by *SFN*) in tsA201 expressing $Na_V1.5$ or $Na_V1.5$-FLAG. Endogenous α-1-syntrophin was used as a positive control for co-immunoprecipitation with $Na_V1.5$, and α-actin as a negative control.

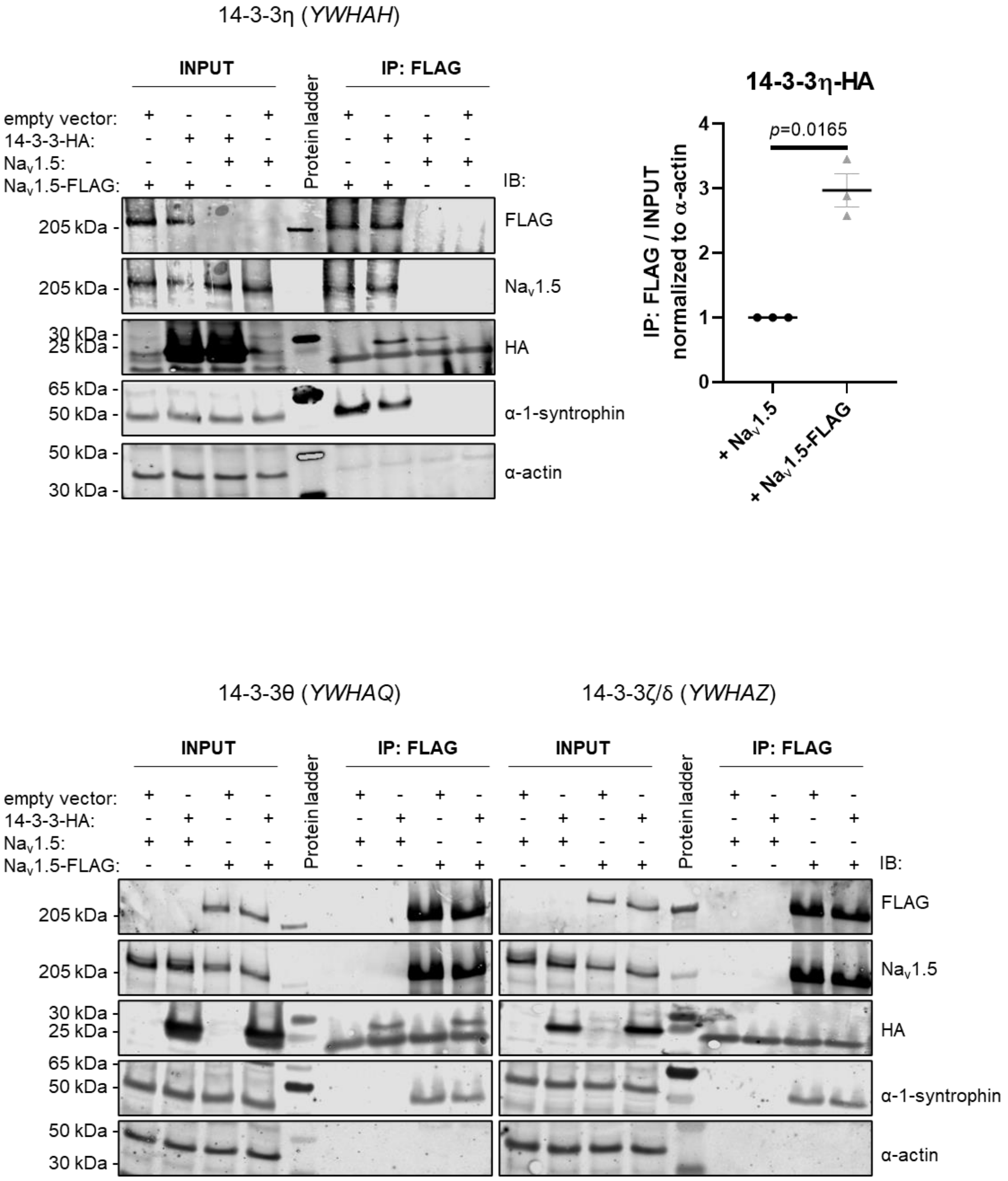


**Supplemental figure S2. Co-immunoprecipitation between Nav1.5 and 14-3-3 isoforms:** 48 hours after transient overexpression of 14-3-3η (encoded by *YWHAH*), 14-3-3θ (encoded by *YWHAQ*) and 14-3-3ζ/δ (encoded by *YWHAZ*) in tsA201 expressing $Na_V1.5$ or Nav1.5-FLAG. Endogenous α-1-syntrophin was used as a positive control for co-immunoprecipitation with $Na_V1.5$, and α-actin as a negative control. Intensity of $Na_V1.5$-immunoprecipitated 14-3-3η-HA was normalized to the intensity of the total 14-3-3η-HA divided by the intensity of α-actin. Data are presented as mean ± SEM from three biological replicates and are normalized to the control condition ("+ $Na_V1.5$"). Individual *p*-value, calculated with one-sample two-tailed t-test with hypothetical mean value=1, is indicated in the panel.

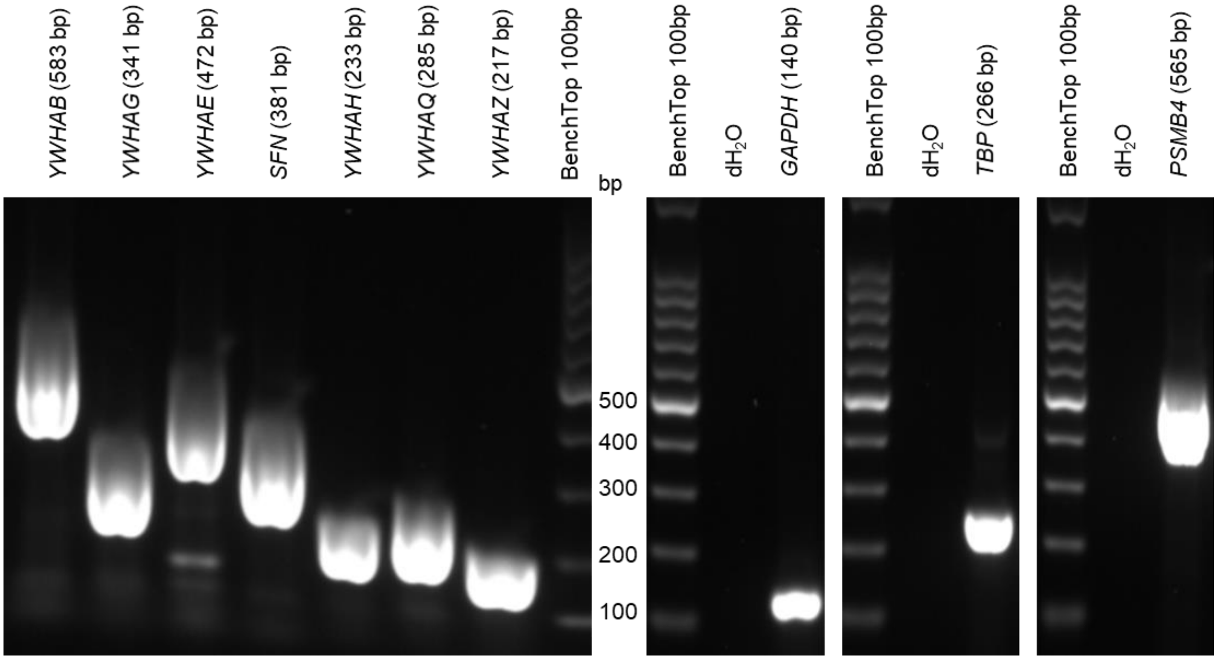


**Supplemental figure S3. Endogenous expression of 14-3-3 isoforms in tsA201 cell line.** Conventional RT-PCR on tsA201 WT cells showing endogenous transcripts of all seven 14-3-3 isoforms (14-3-3β/α encoded by *YWHAB*, 14-3-3γ encoded by *YWHAG*, 14-3-3ε encoded by *YWHAE*, 14-3-3σ encoded by *SFN*, 14-3-3η encoded by *YWHAH*, 14-3-3θ encoded by *YWHAQ*, and 14-3-3ζ/δ encoded by *YWHAZ*) as well as housekeeping genes (glyceraldehyde-3-phosphate dehydrogenase encoded by *GAPDH*, TATA-binding protein encoded by *TBP*, and proteasome 20S subunit β4 encoded by *PSMB4*) with their expected amplicon sizes.

a

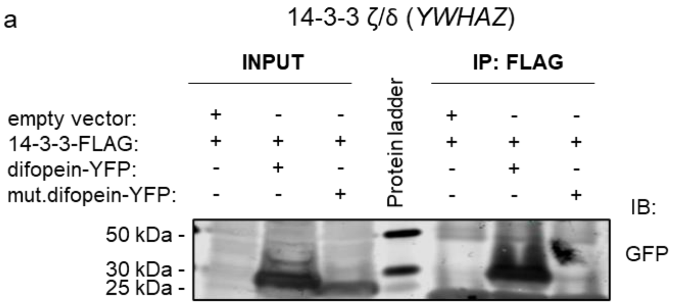


b

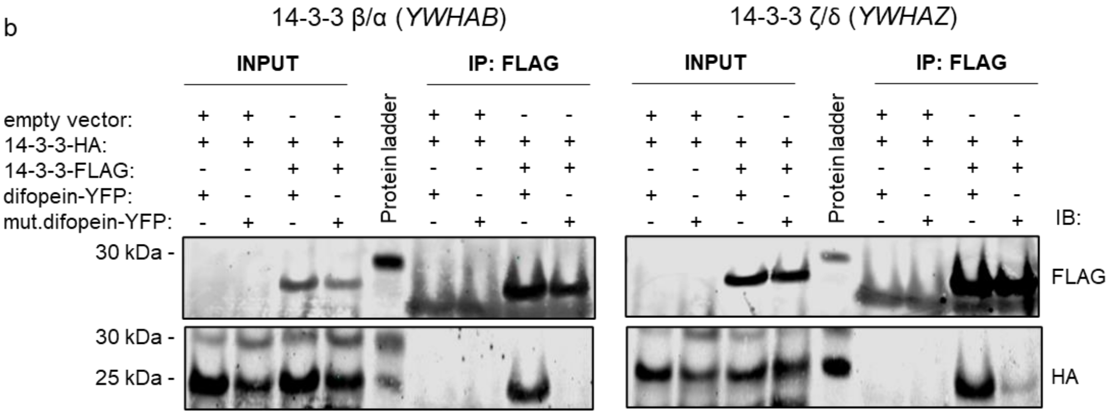


**Supplemental figure S4. Validation of difopein as an efficient competitor for 14-3-3/ligand interaction.** Co-immunoprecipitation analysis was performed 48 hours after transient overexpression of 14-3-3 in tsA201 WT cells. **(a)** Difopein strongly co-immunoprecipitated with 14-3-3 proteins, while the mutant of difopein did not. YFP-tagged difopein and its mutant were revealed with an anti-GFP antibody. **(b)** Difopein stabilized 14-3-3 dimers occupying its binding grooves, preventing 14-3-3 interactions with other ligands. The mutant of difopein did not stabilize 14-3-3 dimers and hence could be used as the closest relevant control for experiments involving difopein.
